# Risk estimates for microcephaly related to Zika virus infection - from French Polynesia to Bahia, Brazil

**DOI:** 10.1101/051060

**Authors:** Michael A Johansson, Luis Mier-y-Teran-Romero, Jennita Reefhuis, Suzanne M Gilboa, Susan L Hills

## Abstract

Zika virus (ZIKV) infection during pregnancy has been linked to birth defects,^1^ yet the magnitude of risk remains uncertain. A study of the Zika outbreak in French Polynesia estimated that the risk of microcephaly due to ZIKV infection in the first trimester of pregnancy was 0.95% (95% confidence interval: 0.34-1.91%), based on eight microcephaly cases identified retrospectively in a population of approximately 270,000 people with an estimated 66% ZIKV infection rate.^2^

---

In the current outbreak, thousands of suspected cases of infants with microcephaly or other central nervous system developmental anomalies possibly associated with ZIKV infection have been reported in Brazil. To estimate the magnitude of microcephaly risk in Brazil, we analyzed data from Bahia (Fig. A).^3^ Serosurvey data from Yap^4^ and French Polynesia^2^ indicate that reported Zika cases only represent a small fraction of the number of ZIKV infections that actually occur. The infection rate in Bahia cannot be reliably inferred from currently available data, so we assumed that it could range from 10% to 80% based on estimates from Yap and French Polynesia (66-73%)^2,4^ and reports from non-outbreak ZIKV serosurveys (6-40%, Supplementary Materials). We apportioned this risk across 2015 according to the temporal distribution of reported cases (Fig. B) and assessed the association of infection risk with microcephaly cases reported in the Brazilian Live Births Information System^5^ between July 2015 and February 2016 (as of March 21, 2016, accounting for a reporting delay, Fig. C).

**Figure.**
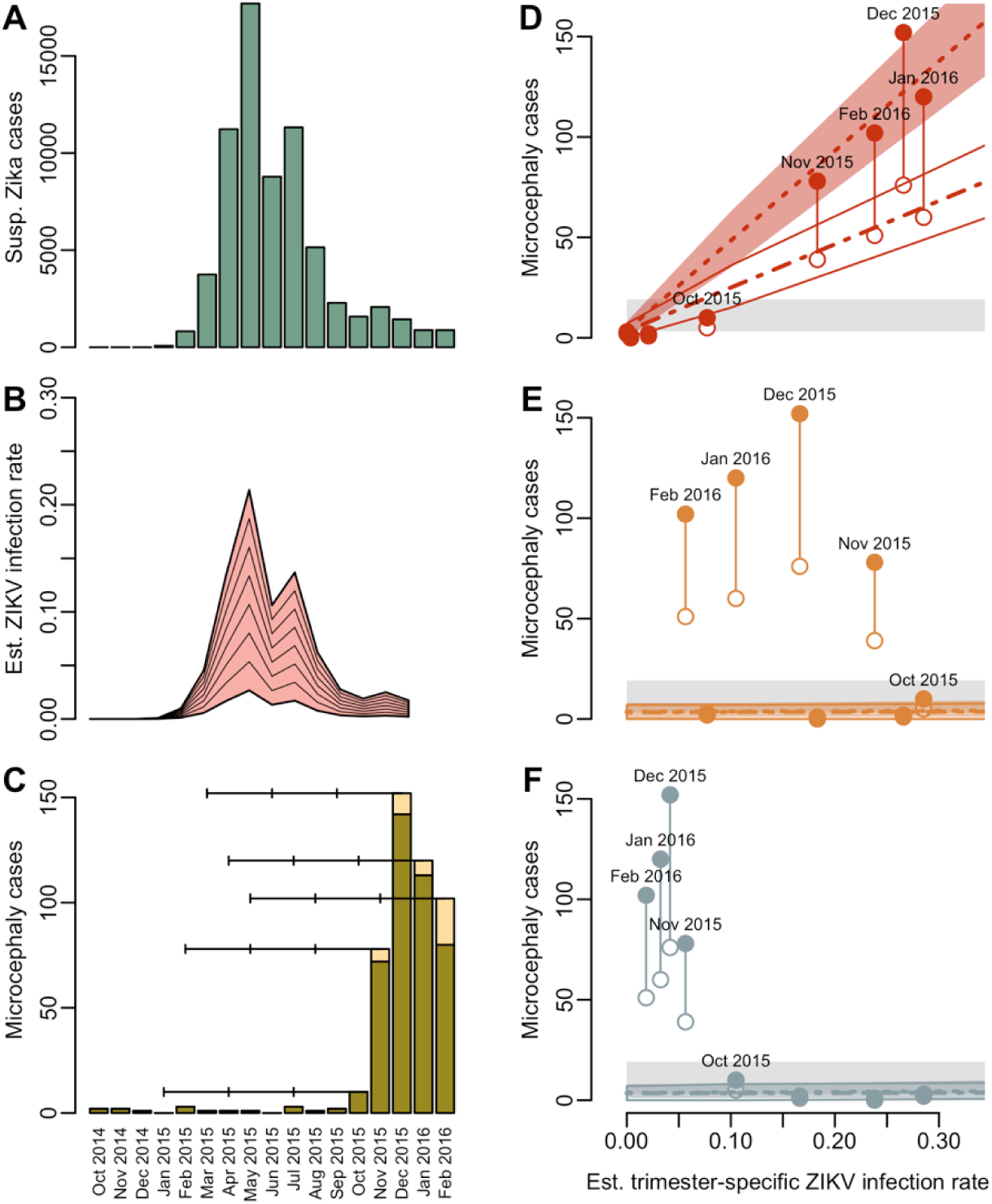
The relationship between trimester-specific ZIKV infection risk and microcephaly in Bahia, Brazil. (**A**) The approximate number of suspect Zika cases reported in Bahia by month^3^. (**B**) The estimated ZIKV infection rate, assuming an overall infection rate of 10-80% (lines in 10% increments). (**C**) Microcephaly cases in Bahia including reported cases (dark green)^5^ and estimated additional cases (yellow), accounting for reporting delays (Supplementary Material). Horizontal lines indicate the approximate gestational period by trimester for pregnancies coming to term in October 2015 through February 2016. (**D, E, F**) The solid points represent the total number of microcephaly cases for each birth cohort (July 2015-February 2016) in Bahia (adjusted for reporting delays) relative to the estimated infection rate for the first (**D**), second (**E**), and third (**F**) trimesters if the overall infection rate for 2015 was 50% (other overall infection rates produced qualitatively similar results). The open points represent 50% of this value (reflecting potential over-reporting) and the grey area represents expected baseline microcephaly rates of 2-12 cases per 10,000 births. Model-fitted estimates and 95% credible intervals for microcephaly cases across trimesters are shown for data with (dot-dashed) and without over-reporting (dashed). For the second and third trimesters the estimates are negligible and overlap.

Considering different infection rate scenarios (from 10% to 80%), possible over-reporting (0% or 100%), and an uncertain baseline microcephaly rate (2-12 cases per 10,000 births), microcephaly risk was strongly associated with infection risk in the first trimester, with a negligible association in the second and third trimesters, consistent with the association found in population-level estimates for French Polynesia^2^ (Supplementary Material). Estimated baseline microcephaly risk was low, approximately 2 per 10,000 births (Tables S1–2, Fig. D–F), but estimated risk due to infection in the first trimester ranged from 0.88% (95% Credible Interval (CI): 0.80-0.97%) with an 80% overall ZIKV infection rate and 100% over-reporting of microcephaly cases to 13.2% (95% CI: 12.0-14.4) with a 10% ZIKV infection rate and no over-reporting. The lower end of this range is similar to the approximately 1% risk estimated for French Polynesia, especially if infection rates in Bahia were high (40% or more with over-reporting, 70% or more without over-reporting). It is also possible that the French Polynesia estimate is an underestimate; it is from a single outbreak relying on retrospective microcephaly case detection. Furthermore, higher microcephaly risks have been documented for some other viruses.^2^ Both estimates are consistent with the lack of reported microcephaly cases in Yap; if microcephaly risk due to first trimester ZIKV infection was 0.88-13.2%, 0-4 microcephaly cases would have been expected.

There are uncertainties and limitations to all current estimates of microcephaly risk associated with ZIKV infection. First, available data are very limited, especially in recently affected areas like Bahia where infection rates are unknown and microcephaly cases are still being reported and evaluated. The limited information on ZIKV infection rates is compounded by difficulty in clinical confirmation of microcephaly, as evidenced by low confirmation rates in the independent, temporary microcephaly reporting system established by Brazil in late 2015.^6^ Carefully designed serosurveys and data from other locations can help refine these estimates.

Recent studies show associations between symptomatic ZIKV infection during all trimesters and adverse pregnancy outcomes^7^ and potential peak risk during gestational weeks 14-17.^8^ How these outcomes relate to the clear association between first trimester risk and microcephaly at the population level in French Polynesia and Bahia is unclear. On the population level, the temporal relationship is confounded by varying infection risk, gestational ages, and fetal outcome assessment. Meanwhile in clinically described cases, infections in early pregnancy may be more difficult to document as pregnancy status may not be known and outcomes associated with mild or asymptomatic ZIKV infection during pregnancy have yet to be described. Risk of adverse events may be higher in symptomatic infections^7^, but mild infections are likely more common and thus may also contribute substantially to the overall burden. Furthermore, microcephaly is only one possible adverse outcome among a spectrum of conditions that may be part of congenital Zika syndrome. A population-level increase in central nervous system anomalies has been observed in both French Polynesia^9^ and Brazil.^6^ More data are needed to refine gestational age-specific risk estimates for microcephaly and these other outcomes related to ZIKV infection.

While much remains unknown about the effects of ZIKV infection during pregnancy, there is a clear association between first trimester ZIKV infection and microcephaly risk from population-level data in French Polynesia and Bahia. If the risk of infection and adverse outcomes is similar in the other areas where ZIKV has spread, many more cases of microcephaly and other adverse outcomes are likely to occur. In light of the growing evidence, it is prudent to take precautions to avoid ZIKV infection during pregnancy^10^ and for health care systems to prepare for an increased burden of adverse pregnancy outcomes in the coming years.

## Acknowledgements

The findings and conclusions in this report are those of the authors and do not necessarily represent the official position of the Centers for Disease Control and Prevention. All data are publicly available and code is available from MAJ upon request. MAJ, LMTR, JR, SMG, and SLH conceived of the research, wrote, and approved of the manuscript. MAJ and LMTR performed the analyses. MAJ received partial support from the Models of Infectious Disease Agent Study program (Cooperative Agreement 1U54GM088558).

## Supplementary Materials

**Materials and methods**

**Pregnancy data**. Over the years 2010-2014, approximately 16,000 live births (range: 13,141-20,713 births) occurred in Bahia per month^5^. To estimate the number of microcephaly cases expected in the area of Yap where the first Zika outbreak was described, we used a birth rate estimate of 16.5 births per 1,000 people (Federated States of Micronesia)^11^ and a population size of 7,391 people^4^.

**Microcephaly data**. Monthly microcephaly data were collected from the Live Births Information System (SINASC)^5,12^. Recent data reflect a reporting lag; the most recent months have many fewer reported births than expected and those numbers increase in subsequent months as additional records are entered retrospectively. We accounted for this lag by calculating the proportion of microcephaly cases among all reported births in a given month and multiplying this by 16,000 to estimate the total number of microcephaly cases expected in a full birth cohort of approximately 16,000 births.

We accounted for uncertainty in reporting and baseline microcephaly rates using data from other surveillance systems. As of March 19, 2016, the independent microcephaly surveillance system in Brazil^6,12^ reported that 59% of fully investigated cases of microcephaly or central nervous system anomalies in Bahia had signs of congenital infection due to ZIKV or other infections. Across all of Northeastern Brazil, the confirmation rate was lower, 44%. In contrast to this new surveillance system, the SINASC data come from an established system with a strict World Health Organization microcephaly case definition. We therefore tested two alternative assumptions for microcephaly cases reported by SINASC: no over-reporting or 100% over-reporting (indicating a 50% confirmation rate). Other causes of microcephaly account for 2 to 12 cases per 10,000 births in the United States^13^, which we considered as the baseline range for microcephaly in Bahia.

**Zika data**. Suspect Zika case data were collected from the Bahia State Secretary of Health Arbovirus Situation Report^3^ using WebPlotDigitizer (arohatgi.info/WebPlotDigitizer). The digitized data were scaled to match overall reported case numbers for 2015. Few suspect cases were tested, however the abundance of rash with relatively little fever and the shape of the epidemic curve which did not parallel reports of suspected dengue cases^14^, suggest that a majority of disease among the reported cases might have been related to ZIKV infection.

There are few population-based estimates of ZIKV infection rates from other outbreaks: on Yap, 73% (95% confidence interval: 68-77%)^4^ and French Polynesia, 66-86% ^2,15^. Seroprevalence estimates for ZIKV between 6% and 40% have been reported, but these reports were not related to specific outbreaks and may have been confounded by the presence of antibodies to other flavivirus ^16^^−^^21^. Infection rates following Zika outbreaks might be similar to rates following chikungunya outbreaks as chikungunya virus is transmitted by the same mosquitoes and has also caused outbreaks in previously naïve populations. Chikungunya outbreaks have resulted in infection rates of 40-60% (range: 17% to 80%) ^22^^−^^27^. As estimation of microcephaly risk is intrinsically related and sensitive to the overall infection risk, we assessed microcephaly risk at a range of population-level ZIKV infections risks, from 10% to 80%. For each infection rate scenario (10% to 80%), we calculated an estimated infection rate per month of 2015 based on the number of reported suspect Zika cases.

**Estimation of microcephaly risk**. We calculated the risk of ZIKV infection during each trimester *p*_*Zt,i*_, where *t* indicates the trimester and *i* indicates the cohort, assuming that the average trimester for a given birth cohort begins in the middle of one month and ends in the middle of another. For example, ZIKV infection risk for the third trimester in the October birth cohort was the sum of half the risk in July, all of the risk in August and September, and half the risk in October. To account for immunity arising from infections occurring before pregnancy or in earlier trimesters, we also estimated the risk of previous ZIKV infection for each cohort at each trimester *p*_*It,i*_. We summed the risk in all previous months including half the risk of the first month of the trimester. We defined the trimester specific microcephaly risk due to ZIKV infection as *p*_*M|Zt*_ and baseline microcephaly risk as *p*_*M0*_, such that the total risk of microcephaly in a given cohort is the sum of baseline risk plus risk in each trimester (discounting for possible previous infection and baseline microcephaly risk):

*P*_*Mi*_ = *P*_*M0*_ + *P*_*M|Z1,i*_(1 − *P*_*I1,i*_)(1 − *P*_*M0*_) + *P*_*M|Z2*_*P*_*Z2,i*_(1 − *P*_*I2,i*_)(1 − *P*_*M0*_) + *P*_*M|Z3*_*P*_*Z3,i*_(1 − *P*_*I3,i*_)(1 − *P*_*M0*_).

This risk is the probability of microcephaly for each cohort of 16,000 births such that the observed number of microcephaly cases in a cohort is *M*_*i*_:

*M*_*i*_ ~ Binomial(*αP*_*Mi*_, 16,000),

where *α* is an over-reporting factor (1 or 2 in our analysis to reflect no over-reporting and 100% over-reporting, respectively). We fit this model in R (R-project.org) using Markov Chain Monte Carlo sampling in JAGS (mcmc-<u>jags.sourceforge.net</u>) via rjags (CRAN.R-project.org/package=rjags). We assigned each *P*_*M|Z*_ parameter a naïve Beta prior and used uniform priors for *P*_*M0*_ (2-12 cases per 10,000 births). Code is available from the authors upon request.

**Supplementary Table 1.** Estimated microcephaly risk by trimester of ZIKV infection, Bahia, Brazil, July 2015-February 2016 with no over-reporting.

| Overall ZIKV infection rate | Estimated microcephaly risk (%) |  |  |  |  |  |  |  |
| --- | --- | --- | --- | --- | --- | --- | --- | --- |
|  | Baseline |  | With 1<sup>st</sup> trimester ZIKV infection |  | With 2<sup>nd</sup> trimester ZIKV infection |  | With 3<sup>rd</sup> trimester ZIKV infection |  |
|  | Mean | 95% CI | Mean | 95% CI | Mean | 95% CI | Mean | 95% CI |
| 10% | 0.023 | (0.02, 0.029) | 13.2 | (12.0, 14.4) | 0.055 | (0.001, 0.2) | 0.063 | (0.002, 0.222) |
| 20% | 0.022 | (0.02, 0.029) | 6.69 | (6.07, 7.35) | 0.028 | (0.001, 0.101) | 0.033 | (0.001, 0.115) |
| 30% | 0.023 | (0.02, 0.029) | 4.52 | (4.10, 4.96) | 0.018 | (0, 0.066) | 0.023 | (0.001, 0.081) |
| 40% | 0.022 | (0.02, 0.029) | 3.44 | (3.12, 3.77) | 0.014 | (0, 0.05) | 0.018 | (0, 0.065) |
| 50% | 0.022 | (0.02, 0.029) | 2.79 | (2.53, 3.06) | 0.011 | (0, 0.039) | 0.015 | (0, 0.054) |
| 60% | 0.023 | (0.02, 0.029) | 2.36 | (2.15, 2.59) | 0.009 | (0, 0.033) | 0.013 | (0, 0.048) |
| 70% | 0.023 | (0.02, 0.029) | 2.05 | (1.86, 2.25) | 0.008 | (0, 0.028) | 0.012 | (0, 0.043) |
| 80% | 0.023 | (0.02, 0.029) | 1.83 | (1.65, 2.00) | 0.007 | (0, 0.025) | 0.011 | (0, 0.04) |

**Supplementary Table 2.** Estimated microcephaly risk by trimester of ZIKV infection, Bahia, Brazil, July 2015-February 2016 with 100% over-reporting.

| Overall ZIKV infection rate | Estimated microcephaly risk (%) |  |  |  |  |  |  |  |
| --- | --- | --- | --- | --- | --- | --- | --- | --- |
|  | Baseline |  | With 1<sup>st</sup> trimester ZIKV infection |  | With 2<sup>nd</sup> trimester ZIKV infection |  | With 3<sup>rd</sup> trimester ZIKV infection |  |
|  | Mean | 95% CI | Mean | 95% CI | Mean | 95% CI | Mean | 95% CI |
| 10% | 0.021 | (0.02, 0.024) | 6.38 | (5.75, 7.01) | 0.025 | (0.001, 0.092) | 0.027 | (0.001, 0.098) |
| 20% | 0.021 | (0.02, 0.024) | 3.23 | (2.92, 3.56) | 0.013 | (0, 0.047) | 0.014 | (0, 0.052) |
| 30% | 0.021 | (0.02, 0.024) | 2.19 | (1.98, 2.41) | 0.009 | (0, 0.031) | 0.01 | (0, 0.036) |
| 40% | 0.021 | (0.02, 0.024) | 1.66 | (1.51, 1.83) | 0.006 | (0, 0.023) | 0.008 | (0, 0.028) |
| 50% | 0.021 | (0.02, 0.024) | 1.35 | (1.22, 1.49) | 0.005 | (0, 0.018) | 0.006 | (0, 0.023) |
| 60% | 0.021 | (0.02, 0.024) | 1.14 | (1.03, 1.25) | 0.004 | (0, 0.015) | 0.006 | (0, 0.021) |
| 70% | 0.021 | (0.02, 0.024) | 0.993 | (0.899, 1.09) | 0.004 | (0, 0.013) | 0.005 | (0, 0.018) |
| 80% | 0.021 | (0.02, 0.024) | 0.883 | (0.798, 0.971) | 0.003 | (0, 0.012) | 0.005 | (0, 0.017) |

